# Once delayed non-invasive remote ischemic preconditioning protects against early stroke by modulating neuroinflammatory responses in rats

**DOI:** 10.1101/2020.05.14.095810

**Authors:** Xiangnan Du, Jian Yang, Yanlong Zhao, Xuemei Wang, Xiaokun Geng

## Abstract

Once delayed non-invasive remote ischemic preconditioning (RIPC) has been proven to provide endogenous protection against injury induced by ischemia–reperfusion in the central nervous system. However, for thus ischemic preconditioning method, it is still unclear how long this protection can maintain and what the underlying mechanism is. In this study, we tested the hypothesis that once delayed non-invasive RIPC protects brain injury at short reperfusion time. The rat was stimulated by transient middle cerebral artery occlusion (MCAo) for 90 min, and subsequent reperfusion was performed at 6 h, 72 h and 7 days after MCAo. RIPC was conducted in both hind limbs 24 h before MCAo for 3 cycles (10 min ischemia/ 10 min reperfusion). The infarct size was measured by 2, 3, 5-triphenyl-2H-tetrazolium chloride (TTC) staining and Cresyl violet (CV) staining. The mRNA and protein levels of inflammatory cytokines in the brain were measured by real-time RT-PCR and ELISA. The results showed that once delayed non-invasive RIPC reduced the infarct size, improved neurological functions and behavioral performance at 6 and 72 h post-stroke. There was no change by reperfusion at 7 d after MCAo. RIPC reduced the levels of TNFα, IL-1β and IL-6 in the brain at 72 h post stroke. It also reduced the levels of TNFα and IL-1β when reperfusion at 6 h after MCAo. Our results strongly supported that once delayed non-invasive RIPC protects against stroke as a non-invasive neuroprotective strategy, which maintained for both short and middle term ischemic reperfusion time. The protective effect is mediated by the modulation of inflammatory response in the ischemic brain.

## Introduction

Ischemic stroke is considered to be the third most fatal and disabling disease in the world. At present, the most effective treatment for stroke is intravenous thrombolysis or intravascular interventional treatment within several hours after the onset of stroke. Unfortunately, the proportion of patients who can be treated within a few hours is less than 5%, and even if the infarction is lifted and blood reperfusion is established, the ischemia-reperfusion injury of the brain tissue cannot be ignored, which causes the current unsatisfactory results. Neuroprotective drugs developed over the years have been proven effective in animal models of stroke, but have poor clinical efficacy [1]. Therefore, it is urgent to find auxiliary or alternative treatment to further improve the treatment effect of stroke. In recent years, a variety of remote ischemic preconditioning (RIPC) methods have been tested as feasible treatment strategies for stroke. Our previous studies and other researches have proved that RIPC has a protective effect on stroke in both basic research [2–4] and clinical experiments [5–7]. RIPC is easy to handle and relatively resistant to reperfusion injury, so it has great clinical advantages. However, for the once delayed non-invasive RIPC method, the duration of protection and mechanisms are still unclear.

Preconditioning is a phenomenon in which the brain protects itself against future injury by adapting to low doses of noxious insults [8]. The concept of cerebral ischemic tolerance was first introduced in the early 1990s. Kitagawa et al. reported the neuroprotective effects against neuronal cell death when adding 2 minutes of transient ischemia 24 hours before global cerebral ischemia in rats [9]. As ischemic conditioning is difficult to realize in the *in situ* organ, the concept of RIPC is proposed. RIPC is an endogenous protective mechanism through which the short-term sub-lethal ischemia of remote organs can protect the main organs from further severe ischemia. Acute and delayed preconditioning in both heart and brain has different mechanisms. Acute/early preconditioning performed 1 to 3 hours before stroke onset is related to a rapid response such as changes in ion channel permeability and post-translational modifications of proteins and the protection lasted only several hours. While delayed preconditioning induced 1 to 7 days before stroke onset induced gene activation and protein synthesis and the protection lasted several days [10–13]. In most experiments, the protective effects on the brain need hours and sometimes days to fully manifest; thus, delayed preconditioning has been studied as a more effective strategy. To date, studies on the mechanisms of both cardiac and cerebral preconditioning at the molecular, cellular and tissue levels span nearly 30 years [14]. Many studies have shown that the neuroprotective mechanisms of RIPC by a complex cellular regulatory process that involves multiple cellular signaling pathways and leads to enhanced tolerance to ischemia/hypoxia.

So far, there are many ways of RIPC, and their effects are not completely consistent. Previous study showed that non-invasive RIPC 5 min ischemia/ 5 min reperfusion for 3 cycles contributed neuroprotection by activating adenosine A1 receptor [15]. Another study showed that 4 cycles of RIPC (5 min/cycle, 40 min total) decreased the expression of neuroinflammation by improving the peripheral immune cell response [2]. Our previous research showed that 3 cycles of RIPC (10 min ischemia/ 10 min reperfusion) significantly reduced infarct size and neuroinflammation by modulating the expression of HIF-1α [4]. Moreover, studies showed that acute (15 min before ischemia), delayed (24 h before ischemia) and chronic (9 d repeated ischemic conditioning) ischemic preconditioning all reduced infarct size in heart following myocardial ischemia [16]. For the 3 cycles of once delayed non-invasive RIPC (10 min ischemia/ 10 min reperfusion), it has been proven effective in our previous study, whereas, the duration of this protection lasting is still unclear.

In this study, we aimed to provide insights into demonstrating how long the protective effect of once delayed non-invasive RIPC can be maintained. First, we tested the infarct size, neurological and behavioral deficiencies at different reperfusion time compared with MCAo and RIPC+MCAo group. Then, we tested the mRNA and protein levels of pro-inflammatory cytokines in the ischemic brain by real-time RT-PCR and ELISA respectively.

## Materials and methods

Male Sprague-Dawley (SD) rats were purchased from Vital River Laboratory Animal Technology Co., Ltd (Beijing, China). The weights were 280-320g. They are placed in a control room at a temperature of 24 ° C and a standard 12-hour light-dark cycle. They can freely obtain food and water and are randomly divided into different groups. The number of animals per group is 12 to 14. All procedures in this study were conducted in accordance with ethical standards, with the Helsinki Declaration, with national and international guidelines, and have been approved by the Authors Institutional Review Board.

### Middle cerebral artery occlusion (MCAo)

In our experiments, rats were anesthetized with 3-5 % isoflurane in 70 % nitrous oxide and 30 % oxygen, and maintained with 1-3 % isoflurane. For the MCAo model, ischemia and reperfusion was established as previously described [4]. Briefly, the common carotid artery (CCA), right internal carotid artery (ICA), and external carotid artery (ECA) were exposed during the procedure. A silicone-coated nylon suture with a diameter of 0.38 ± 0.02 mm was inserted into the ICA; the silicone-coated nylon suture was within 18-20 mm from the ECA bifurcation to block the MCA, and withdrawed after 90 minutes of occlusion to allow MCA to re-open. Rat rectal temperature was maintained at 37±0.5° C during the entire procedure. The rats in the sham group underwent surgery without MCA occlusion. The cerebral blood flow (CBF) during the surgery before and after occlusion were measured by laser Doppler perfusion monitoring with a laser Doppler probe (PeriFlux System 5000, Perimed AB, Sweden) interfaced to a laptop equipped with the PeriSoft data acquisition software (PeriSoft Systems, Inc., Sweden). The blood gas (PaCO_2_, PaO_2_ [mmHg] and pH) and blood sugar (mmol/L) were also examined during the surgery as previously described [17].

### Remote ischemic preconditioning (RIPC)

In our experiments, delayed non-invasive RIPC was used, in contrast to the invasive, direct femoral artery occlusion. Briefly, twenty-four hours before MCAo, RIPC was conducted in both hind limbs of rats anesthetized with 1-3 % isoflurane. Two strip gauze bandages were tied on the two hind limbs simultaneously to occlude blood circulation for 10 minutes, and then released for 10 minutes to allow reperfusion. The occlusion/reperfusion cycle was repeated for 3 times.

### Behavioral testing

Behavioral tests were conducted as described previously [4, 17]. Behavioral tests were performed to assess rats’ neurological function after stroke, including tail hang tests, home cages test and postural reflexes test. All behavioral tests are performed by a person who does not understand the experimental conditions. We trained rats three days before surgery and tested their baseline one day before surgery. All behavioral tests were evaluated before the animals were sacrificed.

For the tail suspension test, hung the tail of the rat about 10 cm from the ground. Stroke rats will turn to the opposite side (left) of the ischemic hemisphere, and the head will rotate more than 90 °. Each rat was hung for no more than 5 seconds, and each rat was hung a total of 20 times. The percentage of head turns was calculated.

Rats usually used their forelimbs to explore the cage margin. For the home cage limb test, we calculated the number of times when rats’ ipsilateral, contralateral, or both forelimbs contacted the cage wall. Instruct the rat to touch the cage wall 20 times. Use the following formula to calculate the percentage of ipsilateral forelimbs used: [ipsilateral + (both /2)] × 100%.

In a postural reflex test, rat was placed on a table. We held its tail in one hand, and pushed its shoulder nearly 20 cm for 3 times with the other hand. Non-ischemic rats grasped the table vigorously during the push and scored zero. Rats that had less resistance and became stiff during the referral process received 1 point. If the rat was not resistant, the score was 2.

We also used the Longa scoring system to measure neurological deficits at different times after reperfusion to assess stroke outcomes. The scores were based on the following criteria: 0 = no defect, 1 = inability to stretch the left front foot, 2 = circle left 3 = Fall to the left, 4 = Unable to walk away and lose consciousness, 5 = Death.

### Infarct size measurement—TTC staining

For 6 and 72 hours reperfusion animals, the infarct area was measured using 2, 3, 5-triphenyl-2H-tetrazolium chloride (TTC) staining. Measure non-ischemic hemispheres and non-ischemic region and calculate infarct area according to the following formula: [(area of the non-ischemic hemisphere - area of the non-ischemic region in the ischemic hemisphere)/area of the non-ischemic hemisphere] × 100%. Detailed protocols have been described previously.

### Infarct size measurement—Cresyl violet staining

For the long-term reperfusion induced by stroke, the infarct volume was measured by Cresyl violet (CV) staining as the TTC staining method does not reflect the infarct size clearly. Animals were anesthetized and transcardially perfused with N.S., followed by 4% paraformaldehyde in 0.1 M PBS. Brains were post fixed for 12 h in 4% paraformaldehyde and dehydrated in 20% and 30% sucrose in PBS, respectively. Brains were frozen and sectioned coronally (30 μm) and pasted on the slides. Slides were rehydrated with 0.1M PBS for 5 min. Then the slides were stained in 0.1% CV solution at 37°C for 10 min and differentiation in 1% glacial acetic acid. The slides were washed twice with ddH_2_O and immersed in 95% ethanol for 2 minutes. Then, they were cleared twice for 5 min with xylene, sealed with neutral gum, and finally observed under microscope.

### Quantitative RT-PCR analysis

To measure TNFα, IL-1β and IL-6 mRNA expression, total RNA was isolated from the ischemic brain, which was collected on ice and stored at −80°C immediately after the animals were euthanized. RNA was extracted with TRIzol reagent (Cat# 15596-026, Life Technologies, California, USA) according to the manufacturer’s instructions. The purified RNA was then reverse-transcribed into cDNA using the Reverse Transcription System (Cat# E6300S, New England BioLabs® Inc., Ipswich, MA, USA). Quantitative RT-PCR analysis of the mRNA level of TNFα (TNFα, F, TGAACTTCGGGGTGATCGGT, TNFα, R, GGCTACGGGCTTGTCACTCG; IL-1β F, CCCAACTGGTACATCAGCACCTCTC, IL-1β R, CTATGTCCCGACCATTGCTG; IL-6, F, GATTGTATGAACAGCGATGATGC, IL-6, R, AGAAACGGAACTCCAGAAGACC) was performed using the SYBR Green Prime Script kit (RR420A, TAKARA). GAPDH (GAPDH F, TTCCTACCCCCAATGTATCCG; GAPDH R, CCACCCTGTTGCTGTAGCCATA) was chosen as the housekeeping gene. The real-time PCR program steps were: 95°C for 5 min, 45 cycles at 95°C for 5 s, 60°C for 5 s, and 72°C for 10 s, followed by 72°C for 1 min.

### ELISA for quantifying pro-inflammatory cytokines

We measured 3 pro-inflammatory cytokines: TNFα, IL-1β and IL-6 using ELISA kit (Expandbio, Beijing, China) at 6, 72 h and 7 d after MCAo. The procedure was conducted according to the manufacturer’s instructions. Briefly, dilute the standard to five gradients according to the instructions, and keep the sample volume of each gradient in 50 μl. After adding samples, incubate the mixture for 30 minutes at 37°C. Washed 5 times, then added 50 microliters of enzyme-labeled reagent, and incubate again at 37°C for 30 minutes. After 5 times of washing, add chromogenic reagents A and B solution for 15 min. Finally, add stop solution and read the OD value at 450 nm.

### Statistical Analysis

Statistical analysis was performed using Prism 5 (GraphPad software, Inc., La Jolla, USA). Results are presented as the means ± SEM. The difference between means was assessed by the Student’s t test (single comparisons) or by one-way ANOVA with Newman-Keuls Multiple Comparison test as a post hoc test (for multiple comparisons). A value of P < 0.05 was considered statistically significant. The number of rats in each group was 12-14.

## Results

### RIPC reduced infarct size at short reperfusion time following stroke

To test whether once non-invasive delayed RIPC was neuroprotective in different reperfusion time including 6 h, 72 h and 7 d. We measured the weight and infarct size by TTC and CV staining. Firstly, we detected the CBF levels, blood sugar and blood gas between groups which were the requirement of stroke model. CBF levels were monitored during MCAo surgery, and there was no difference between groups. It was reduced nearly 78% of baseline during ischemia and reestablished to 80% of baseline following reperfusion (Fig. 1B). At the same time, there were no differences of blood sugar and blood gas between groups, either (Fig. 1C and D). Results showed that RIPC significantly attenuated the weight loss at 72 h post-stroke (Fig. 2D), while RIPC had no effect of the weight loss at other reperfusion time (Fig. 2A and G). TTC staining results showed that RIPC significantly reduced infarct size from 39.26 ± 1.51 to 31.31 ± 1.68 after reperfusion 6 h (Fig. 2B and C). Similarly, RIPC significantly reduced infarct size from 47.11 ± 1.14 to 36.44 ± 1.82 at 72 h post-stroke (Fig. 2E and F). Whereas, CV staining results showed that there was no significant difference of the infarct size receiving RIPC compared with MCAo group at 7 d post-stroke (Fig. 2H and I).

**Figure 1.**
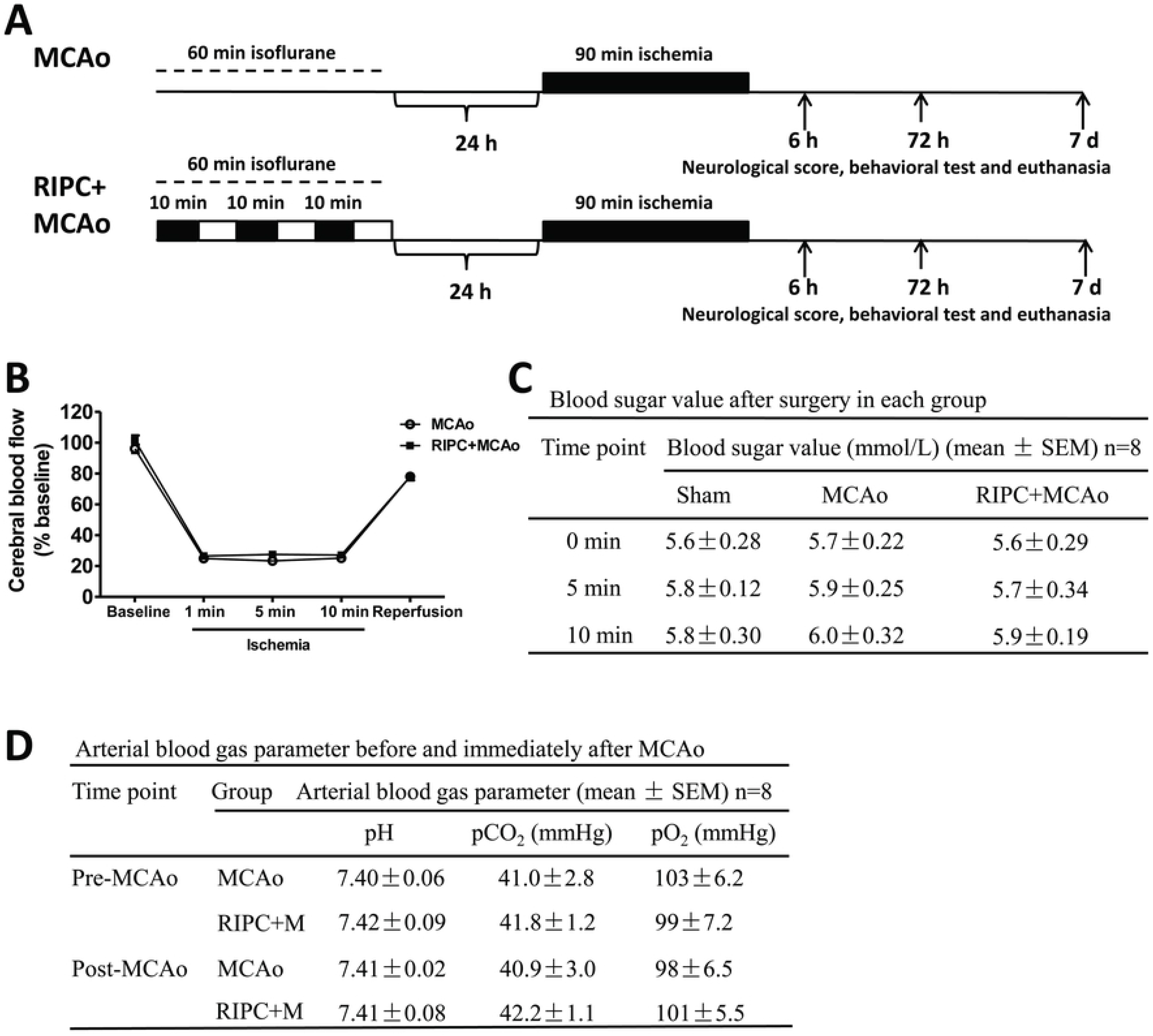
Experimental protocols and model of RIPC. **A.** RIPC was conducted by 3 cycles (60 min total) in both hind limbs under isoflurane. Non-RIPC rats were exposed to the same anesthesia for 60 min. MCAo was induced by 90 min after RIPC 24 h. Neurological score, behavioral test and sample collection at 6 h, 72 h and 7 d post-stroke. **B.** Cerebral blood flow during the MCAo surgery. Cerebral blood flow was measured at five time points, baseline, 1, 5 and 10 minutes of ischemia and reperfusion in the MCAo and RIPC+MCAo groups. Data were normalized to baseline and expressed as percentages. **C.** Blood sugar value after surgery in each group. **D.** Arterial blood gas parameter before and immediately after MCAo. MCAo, middle cerebral artery occlusion; RIPC, remote ischemic preconditioning.

**Figure 2.**
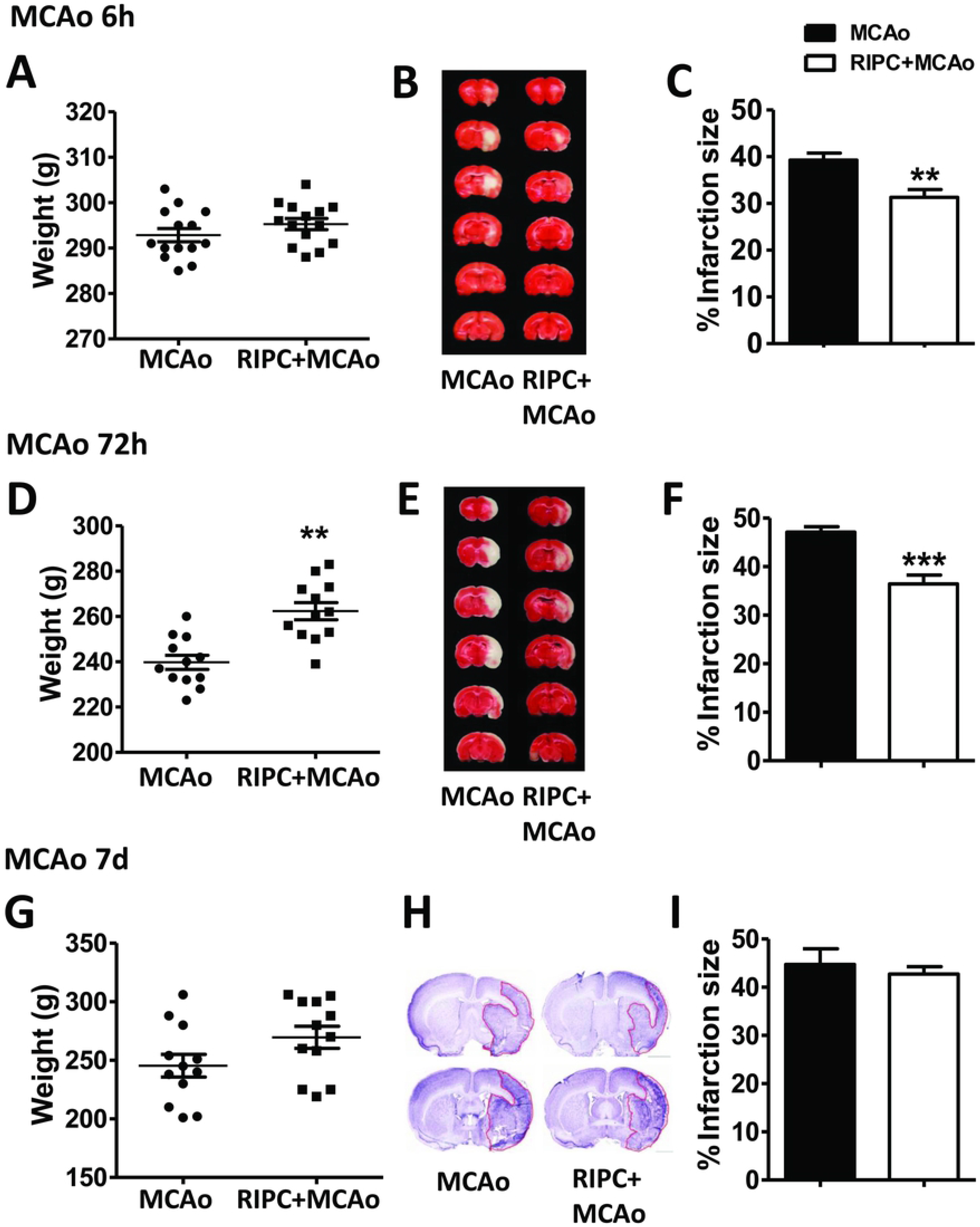
RIPC attenuated the weight loss of rats at 72 h reperfusion post-stroke and reduced infarct size at 6 and 72 h after MCAo. **A, D, G.** Weight of the rats in the MCAo and RIPC+MCAo group. **B, E.** Representative images and infarct volume of TTC staining in the MCAo and RIPC+MCAo group at 6 and 72 h post-stroke. **H**. Representative images and infarct volume of CV staining. **C, F, I.** Statistical analysis of infarct size. Statistical analysis was performed by ANOVA. **, *** p < 0.01, 0.001, vs MCAo, respectively. (N=12-14 per group).

### RIPC improved neurological and behavioral function at early stroke

After validating that once non-invasive delayed RIPC reduced infarct size at acute and middle term ischemic reperfusion time following stroke, we further examined the neurological score and behavioral performance receiving RIPC. The results showed that RIPC significantly attenuated neurological dysfunction at 6 and 72 h post-stroke, while there’s no significant change at reperfusion 7 d after MCAo (Fig. 3A). Such protection of RIPC in these reperfusion time-points were also observed in the behavioral performance test especially in the tail hang test and home cage test, in which RIPC significantly improved the behavioral performance at 6 and 72 h post-stroke (Fig. 3B and C). There was no significant difference in the postural reflex test at any of the reperfusion time-point, but we also observed a decrease trend in RIPC group (Fig. 3D).

**Figure 3.**
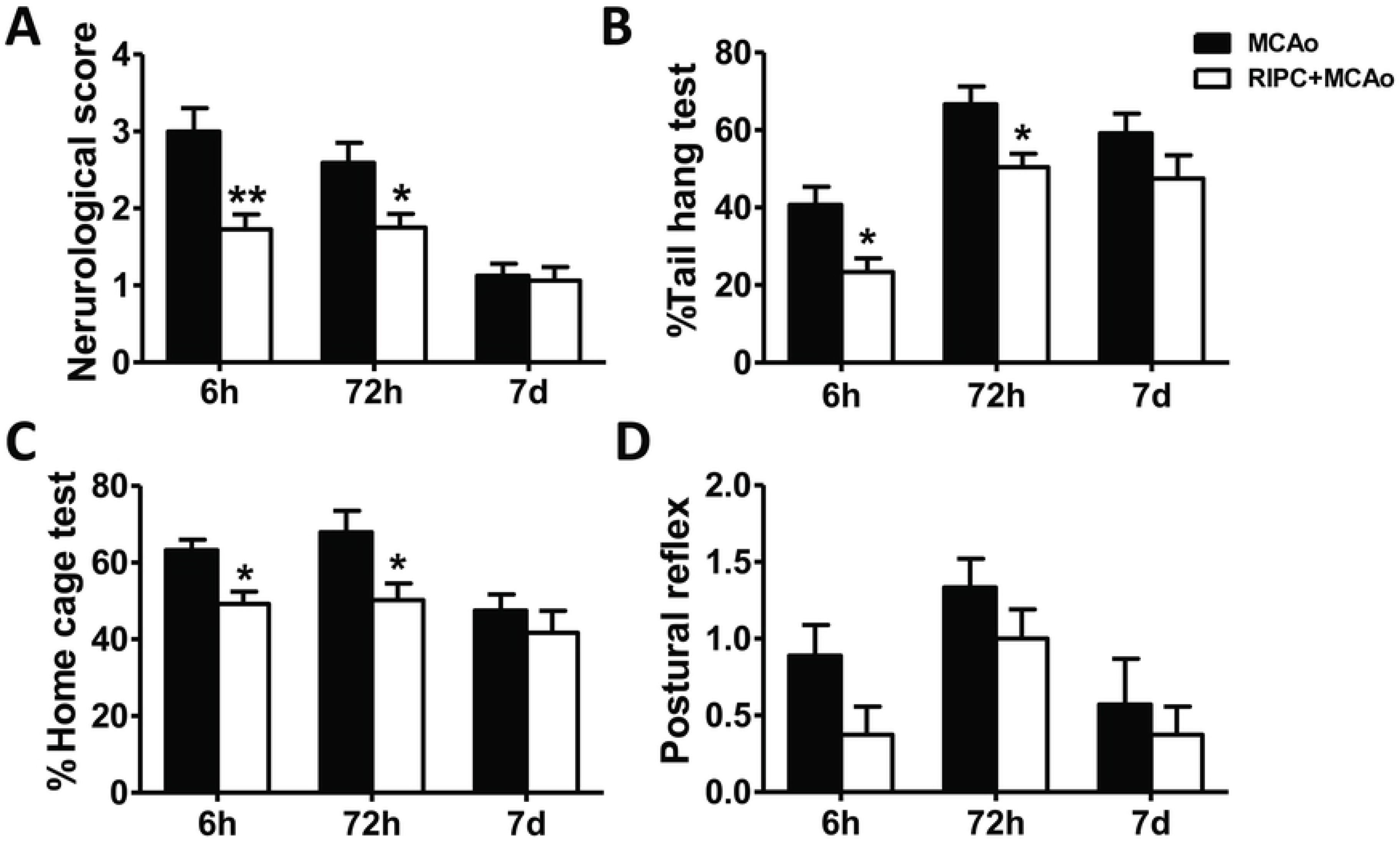
RIPC improved neurological and behavioral function at 6 and 72 h post-stroke. **A.** Neurological score in the MCAo and RIPC+MCAo group at 6 h, 72 h and 7 d post-stroke. **B, C, D.** Behavior tests, including tail hang test, home cage test and postural reflex test. Statistical analysis was performed by ANOVA. *, ** p < 0.05, 0.01 vs MCAo, respectively. (N=12-14 per group).

### RIPC down-regulated pro-inflammatory factors in the ischemic brain at short-term ischemic reperfusion time following stroke

To test how the inflammatory status was regulated by RIPC at different reperfusion time after MCAo, we measured the effect of RIPC on the mRNA and protein levels of the pro-inflammatory cytokines, including TNFα, IL-1β and IL-6 in the ischemic brain. The results showed that MCAo increased the mRNA and protein levels of TNFα in the ischemic brain at any reperfusion time we tested (Fig. 4A and 5A). However, RIPC significantly decreased TNFα expression at 6 and 72 h post-stroke compared with MCAo group at both mRNA and protein levels, while no significant change was observed at 7 d post-stroke (Fig. 4A and 5A). Similar results were observed in IL-1β and IL-6 expression. It is showed that MCAo up-regulated IL-1β mRNA and protein expression at 6 and 72 h after reperfusion and only mRNA level at 7 d after reperfusion, while RIPC decreased the IL-1β expression at mRNA and protein 72 h post-stroke (Fig. 4B and 5B) and mRNA level 6 h post-stroke (Fig. 4B). Whereas, no any significant changes of IL-1β expression were observed at 7 d following MCAo (Fig. 4B and 5B). Finally, it was showed that RIPC down-regulated the expression of IL-6 on the middle-term of reperfusion time (72 h post-stroke) (Fig. 4C and 5C).

**Figure 4.**
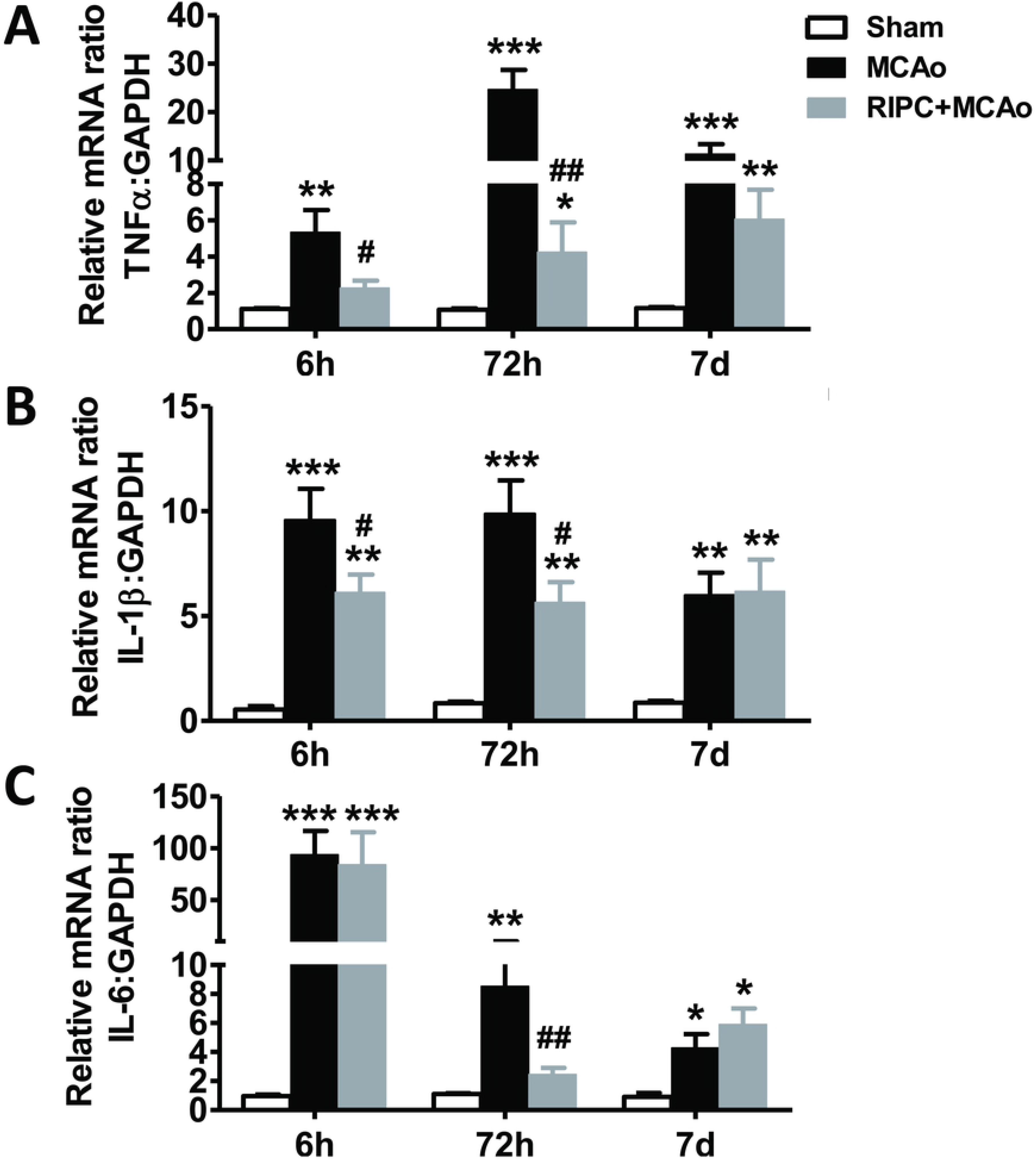
RIPC down-regulated the mRNA levels of pro-inflammatory factors in the ischemic brain. **A, B, C.** The mRNA levels of TNFα, IL-1β and IL-6, respectively. Statistical analysis was performed by ANOVA. *, **, *** p < 0.05, 0.01, 0.001 vs Sham, respectively. #, ## p < 0.05, 0.01 vs MCAo, respectively. (N=12-14 per group).

**Figure 5.**
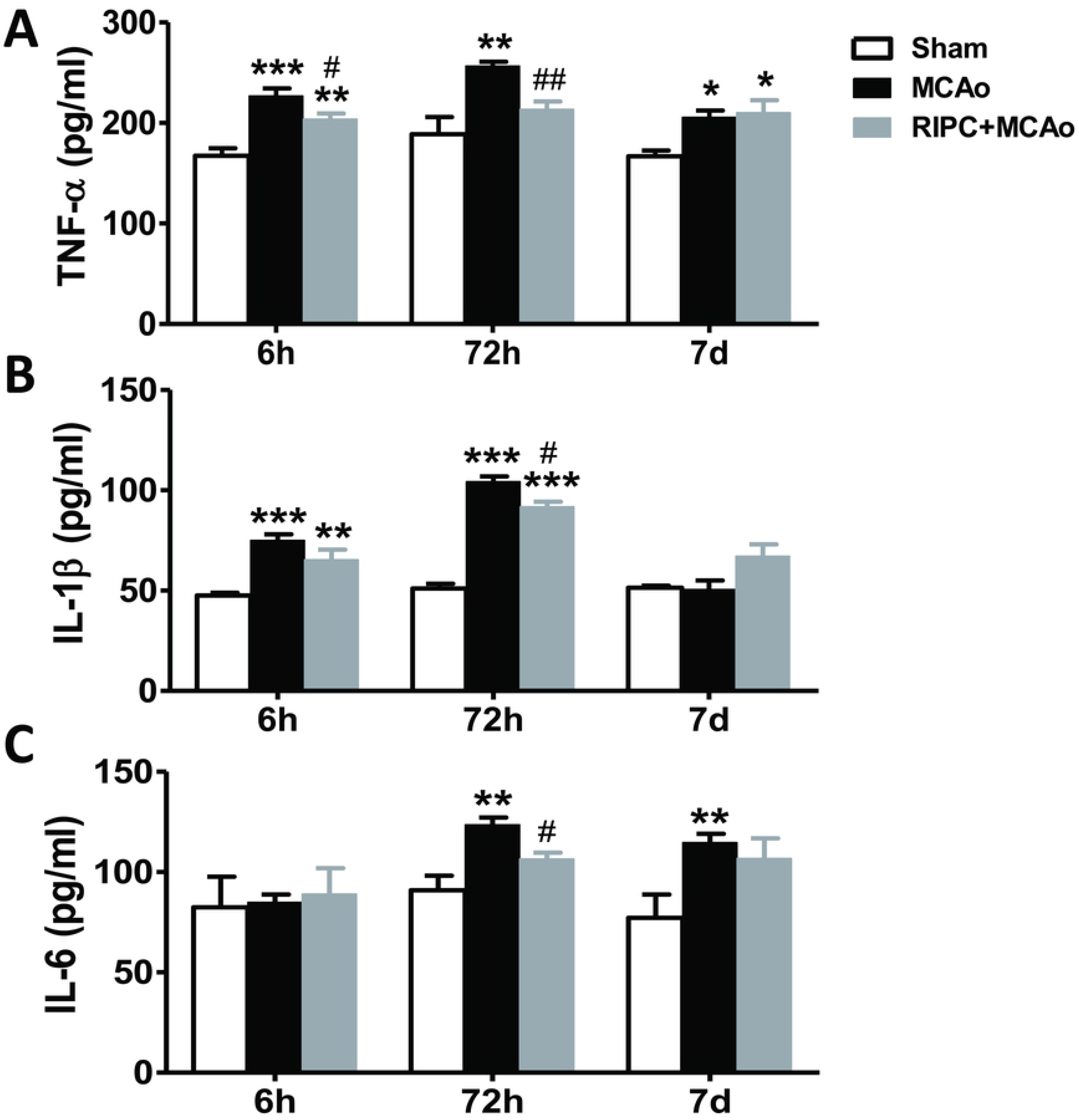
RIPC down-regulated the protein levels of pro-inflammatory factors in the ischemic brain. **A, B, C.** The protein levels of TNFα, IL-1β and IL-6, respectively. Statistical analysis was performed by ANOVA. *, **, *** p < 0.05, 0.01, 0.001vs Sham, respectively. #, ## p < 0.05, 0.01 vs MCAo, respectively. (N=12-14 per group).

## Discussion

In the present study, we investigated the neuroprotection of once delayed non-invasive RIPC against stroke at different reperfusion time. An important finding of this study was that once delayed non-invasive RIPC reduced infarct size, attenuated the loss of neurological function and behavioral performance only at acute reperfusion time - 6 h post-stroke and middle-term reperfusion time - 72 h. However, for the long-term ischemic reperfusion injury, such as 7 d, there was no protection by RIPC. Moreover, RIPC significantly reduced the mRNA and protein levels of pro-inflammatory cytokines including TNFα, IL-1β and IL-6 in the ischemic brain reperfusion 6 and/or 72 h post-stroke. However, there was no difference between MCAo and RIPC+MCAo group on ischemic reperfusion 7 d. Collectively, these findings showed that this ischemic preconditioning method - once delayed non-invasive RIPC protected against stroke as a non-invasive neuroprotective strategy just at short term reperfusion time. The protective effect was mediated by the modulation of inflammatory response in the ischemic brain.

As early as 1986, Murry et al. proposed the concept of ischemic preconditioning with the findings that 4 cycles of a 5 min ischemia/ 5 min reperfusion had a protection in myocardial ischemia [12]. They indicated that ischemic preconditioning was a short period of ischemia, which did not affect the ischemic tissue, but had a protective effect on subsequent, prolonged ischemia. While Kitagawa et al. first introduced cerebral ischemic tolerance in the early 1990s. They discovered 2 min of transient ischemia 24 h before global cerebral ischemia had a neuroprotective effect against neuronal cell death [9, 18]. Researchers regarded ischemic preconditioning as a powerful tool in understanding the endogenous mechanisms by which the ischemic organs are protected [19]. In terms of clinical applicability for myocardial infarction and stroke treatment, RIPC had advantages over conventional ischemic preconditioning by reducing the higher risk directly to the ischemic organ [20, 21]. RIPC refers to a repeated transient ischemia/ reperfusion in a remote organ to prevent prolonged ischemia of other vital organ, which now was widely used in the protection of heart and brain ischemia. In contract to invasive RIPC, we mainly focus on non-invasive method which is often established by tourniquet or strip gauze bandages. The important findings of our study were that once delayed non-invasive RIPC (3 cycles of 10 min ischemia/ 10 min reperfusion) reduced infarct size, attenuated the loss of neurological function and behavioral performance only at acute (6 h) and middle-term (72 h) reperfusion time, but not long-term (7 d) (Figs 2 and 3). The results were consistent with our previous study, in which we have demonstrated that RIPC reduced ischemic/ reperfusion injury at 48 h post-stroke [4]. Moreover, the results were consistent with the data which published by Perez-Pinzon MA et al. They found that ischemic preconditioning *in situ* protected rats against ischemic neuronal damage after 3 but not 7 d of reperfusion following global ischemia [22]. Although different ischemic preconditioning positions were used, these results supported that transient ischemic preconditioning treatment could not maintain long-term protection. However, other studies showed that once rapid non-invasive RIPC (3 cycles of 15 min ischemia/ 15 min reperfusion) had chronic protective effect against distal MCAo even after 60 d [3]. And these contradictory conclusions can be explained by the different model chosen for the two experiments. Moreover, these results supported that RIPC was more effective to improve the infarct in the cortex rather than basal ganglia injury

Inflammatory response plays an important role in the pathogenesis of ischemic stroke. A large number of studies have shown that neuroinflammatory response is involved in the prognosis of cerebral ischemia-reperfusion injury [23, 24]. Therefore, in theory, inhibiting the inflammatory response after stroke can reduce stroke injury and improve the clinical prognosis. Previous studies have shown that RIPC reduced systemic neuroinflammatory response [2–4, 15]. To examine whether reducing infarction was influenced by the elimination of inflammation in the ischemic brain, we then measured the levels of neuroinflammation in the ischemic brain. We found that RIPC reduced the mRNA and protein levels of pro-inflammatory cytokines, including TNFα, IL-1β and IL-6 at acute and middle terms of reperfusion in the ischemic brain, but had no effect on long term reperfusion (Figs 4 and 5). The decrease of cytokines release is consistent with the reduction of the infarct size, suggesting that once delayed non-invasive RIPC improved the regional ischemia by reducing the expression of neuroinflammatory response in the ischemic brain after short and middle term of reperfusion time.

In fact, RIPC has been moved into clinical trials for several years and it has been proven to be effective in the prevention or treatment of cerebrovascular disease [5–7, 20]. The importance of our study was to confirm the limited therapeutic time window of RIPC we used. For different ischemic preconditioning methods and stroke models, RIPC may have different protective effect. Only known that how long it works, can we explore the mechanism and complete further clinical transformation.

There are also several limitations in the present study. First of all, inhibitors of inflammatory cytokines need to be used to demonstrate the interaction between brain injury and inflammatory response in the future study. Secondly, the mechanism of RIPC should be further explored, such as whether the hypothalamus-pituitary-adrenal axis is involved in the protective effect of RIPC. Last but not the least, RIPC was used only once in our experiment, and the parameters of RIPC worth studying to obtain a chronic protective effects for further clinical application.

## Conclusion

In this study, we provided strong evidence that once delayed non-invasive RIPC protects against stroke as a non-invasive neuroprotective strategy which just at short and middle term ischemic reperfusion time. The protective effect was mediated by the modulation of inflammatory response in the ischemic brain.

## Acknowledgments

We thank Xuan Liu and Haiteng Ji for their long-term feeding of experimental animals.

